# Interconduit pit membranes of temperate angiosperms undergo changes in pit membrane thickness and electron density within the final growth ring

**DOI:** 10.64898/2026.07.31.741446

**Authors:** J.R. Schepers, L.M. Silva, C. Carmesin, A. Huppenberge, L. Kaack, N. Korayem, C.L. Trabi, S. Jansen

## Abstract

- Pit membranes play a crucial role in water transport between neighbouring xylem conduits by providing hydraulic safety and flow resistance. While effects of pit membranes thickness on embolism resistance and temporal changes in the ultrastructure of pit membranes have been documented between sapwood and heartwood, these changes are largely unknown within conduits of the current-year.
- We studied interconduit pit membranes of branches of eight angiosperm species by sampling wood from a temperate forest over four consecutive seasons, focusing on the latest growth ring. We quantified the pit membrane thickness and greyscale intensity (as a proxy for electron density) using transmission electron microscopy and image analysis.
- Our observations showed considerable interspecific variation in changes to pit membrane thickness and electron density. Several species exhibited the thinnest pit membranes and highest electron density at the end of a growing season, while others showed minimal variation over time. Intra-tree variation showed that changes in pit membrane shrinkage and electron density were associated with conduit diameter: pit membranes in wide conduits showed larger modification over time than narrow ones.
- Our results indicate seasonal changes in the structure and chemistry of angiosperm pit membranes, even within the latest growth ring. While their shrinkage might increase resistance to flow, the occurrence of darker pit membranes indicates coating and penetration by polar lipids, affecting the behaviour of gas-liquid interfaces. We speculate that the degree of modification that pit membranes undergo is mechanistically driven by the sap flow rate, and possibly conduit dimensions.

## Introduction

The secondary xylem of woody angiosperms provides structural stability of plants and facilitates long-distance transport of xylem sap by conduits. Conduits are specialised xylem cells, such as vessels and tracheids, which transport water and dissolved minerals within plants (Tyree & Zimmermann 2002; Ziemińska 2023). Adjacent conduits are connected via bordered pits, which have pit membranes highly influencing the hydraulic resistance (Zimmermann & Brown 1971; Choat *et al*. 2008). Pit membranes consist of cellulose microfibril aggregates, which develop from hydrolysed primary cell walls in bordered pit pairs of conduits (Schmid & Machado 1968; Catesson 1983; Pereira *et al*. 2018). These microfibrils span the central plane of bordered pit pairs with an average diameter of ca. 5 μm (Kaack *et al*. 2021). Pit membranes have a species-specific thickness between 200 and ca. 1500 nm (Jansen *et al*. 2009; Li *et al*. 2016). Angiosperm pit membranes include mesoporous media between vessels and tracheids, which xylem sap molecules have to cross many times in their xylem pathway from roots to minor leaf veins (Kaack *et al*. 2019). They show a multi-layered nature with a pore volume fraction of about 81% and a low tortuosity (Zhang *et al*. 2020, 2024).

Pit membranes are typically described to play a crucial role in water transport by providing about 50% of the xylem hydraulic resistivity (Choat *et al*. 2008; Kaack *et al*. 2019), with the remaining resistivity provided by the inner conduit walls and perforation plates between individual vessel elements (Ellerby & Ennos 1998; Hacke *et al*. 2006; Christman & Sperry 2010). At the same time, interconduit pit membranes are involved in gas movement from embolised (i.e., gas-filled) conduits to sap-filled ones, contributing most likely to propagation of embolism formation and thus determining embolism resistance (Zimmermann 1983; Sperry *et al*. 1988; Silva, Pfaff, *et al*. 2024; Silva, Pereira, *et al*. 2024). In particular, pit membrane thickness has been associated with embolism resistance (Lens *et al*. 2011; Li *et al*. 2016), given that pit membrane thickness determines the number of pore constrictions within pore pathways (Kaack *et al*. 2021). The narrow pathways in pit membrane pores act as a bottleneck, which determine the movement of fluids and changes in gas-liquid interfaces, such as snap-off events and nanobubble formation (Ingram *et al*. 2023; Silva *et al*. 2025).

During a growing season, cambial activity leads to the formation of a new growth ring of secondary xylem, which is known to reflect seasonal changes and growth conditions (Fritts 1976; Fritts *et al*. 1991; Schweingruber 1996). Vessel dimensions (diameter and length) and frequency as well as wood density may undergo considerable variation, both within a single growth ring, and across consecutive growth rings (Carlquist 1975, 2001; Rathgeber *et al*. 2016; Olson *et al*. 2020). These seasonal changes generally result in a variation of growth ring width as a response to climatic conditions, and can be described as intra-annual density fluctuations (De Micco *et al*. 2016). Pit membranes in angiosperms are known to undergo micromorphological modifications by coating and/or shrinkage, as well as changes in their chemical composition across growth rings, and especially between sapwood and heartwood (Kininmonth 1972; Wheeler 1983; Carmesin *et al*. 2023). These temporal changes in the ultrastructure can be measured with scanning and transmission electron microscopy, allowing to see details of pit membrane structure (Plavcová et al. 2011). Polar lipids coat the inner conduit walls and pit membranes, which become electron dense under transmission electron microscopes (TEM) when osmium is bound to the double carbon bonds of unsaturated fatty acid chains of polar lipids. These amphiphilic lipids have also been referred to as the S3-layer or tertiary wall (Liese 1963; Schenk *et al*. 2017, 2018), and have mainly been identified as phospholipids and galactolipids (Schenk *et al*. 2021; Ingram *et al*. 2023). The polar lipids have been suggested to play a major role in preventing surface bubbles on conduit walls (Šako *et al*. 2026), making these walls more hydrophilic by the hydrophilic heads of galactolipids or phospholipids, while they also provide an essential stability for nanobubbles (Guan *et al*. 2022; Ingram *et al*. 2023). Some of the pioneering studies on pit membrane development and ultrastructure reported that electron density increased with age, while younger developmental stages after hydrolysis showed more transparent and swollen pit membranes (Côté & Day 1962; Harada 1962; Wardrop *et al*. 1963; Schmid & Machado 1968). Further ageing-effects of pit membranes were observed within a growing season for *Vitis vinifera* (Sorek *et al*. 2021), and across growth rings in *Clematis vitalba* (Carmesin *et al*. 2023), suggesting a reduction of pit membrane thickness with age.

Both the time frame considered and the low number of species studied do not allow us to make general conclusions and functional implications about how fast interconduit pit membranes in sapwood undergo changes. While pit membrane shrinkage is reported to occur in plants in the field (Jansen *et al*. 2018) and is known to be largely irreversible (Zhang *et al*. 2017, 2020; Kotowska *et al*. 2020; Sorek *et al*. 2021), observations of pit membrane encrustation or coating can also be temporary (Wheeler 1981; Yamagishi *et al*. 2024). The porosity of pit membranes has been found to decrease considerably when pit membranes become dehydrated, which may cause a shrinkage by ca. 50% compared to fresh, non-dehydrated pit membranes (Zhang *et al*. 2017, 2020; Kotowska *et al*. 2020). This reduced porosity, and the reduction of the pore voids or pore spaces in dried, shrunken pit membranes, is suggested to increase embolism resistance of angiosperms, as has been observed to occur in grapevine (Sorek *et al*. 2021). There are few other studies suggesting an increased embolism resistance during summer drought in other angiosperms (Huang *et al*. 2024; Ma *et al*. 2024). Additionally, polar lipids have been associated with a dynamic surface tension, which play a role in the formation of surfactant-coated surface bubbles, while avoiding embolism (Schenk *et al*. 2018). However, it is unknown if an increase in electron density of pit membranes is associated with embolism resistance. Only few studies have investigated changes in the chemical composition and porous-medium characteristics of pit membranes between conduits, and how these could be potentially associated with functional consequences for water transport. Wheeler (1981) demonstrated that pit membranes of *Fraxinus americana* show a coating during winter, which is largely removed in spring, while Pesacreta et al. (2005) observed a non-microfibrillar coating in overwintering vessels and heartwood of *Triadica sebifera*. Pit membrane encrustations associated with structural and chemical variation were also observed in *Fraxinus mandshurica* before leaf senescence, but disappeared before leaf flushing in spring (Yamagishi et al. 2024).

Here, we aim to investigate the ultrastructure of interconduit pit membranes in eight angiosperm species to record potential changes in their ultrastructure across a full year, including all four seasons. Since long-term structural changes over multiple years can be expected, we explicitly aim to look at potential changes over the course of a single year. Based on earlier records of few species (Schmid & Machado 1968; Sorek *et al*. 2021), we expected that pit membranes would gradually shrink and become more electron dense, thus appearing darker on TEM images. Our specific hypotheses are that: (1) pit membrane thickness declines, and (2) pit membranes become darker from spring to winter.

## Material & Methods

### Plant material and sample preparation

We selected eight temperate angiosperm tree species located around Ulm University and at the Botanical Garden of Ulm University. The selection was based on availability and coverage of a wide range of pit membrane thicknesses (Kaack *et al*. 2021; Guan *et al*. 2022). The species *Acer pseudoplatanus* L.*, Betula pendula* R.*, Carpinus betulus* L.*, Corylus avellana* L.*, Fagus sylvatica* L.*, Liriodendron tulipifera* L.*, Prunus avium* L., and *Tilia cordata* Mill. were investigated. Branches of 0.8 and 1.5 cm in diameter and at least 1 m in length were cut from mature trees at around 1.5 to 3 m height. The cut end was immediately placed in a beaker with water to avoid potential dehydration. Samples were collected across four seasons: summer (August 2019), autumn (November 2019), winter (February 2020), and spring (June 2020; Table S1).

### Transmission electron microscopy

All samples were processed in the lab within two hours at Ulm University following a standard protocol (Kotowska *et al*. 2020; Kaack *et al*. 2021). Small xylem blocks of about 1.5 × 1.5 × 2 mm were cut from thin slivers, including the outer most growth rings and cambium. These blocks were chemically fixed for 24 hours (2.5% glutaraldehyde, 0.1 mol phosphate buffer, 1% sucrose, pH 7.3), washed with PBS buffer, and post-fixed with a 2% osmium tetroxide for one hour. Then, they were gradually dehydrated by submerging them in 50%, 70% and 90% ethanol for 5 minutes each and stained with 20 mg/ml uranyl acetate for 30 minutes. Lastly, the blocks were embedded in Epon epoxy resin (Sigma-Aldrich, Steinheim, Germany). Transverse semithin sections of 500 µm where cut and treated with toluidine blue for light microscopy, and ultrathin sections of 60 nm to 90 nm were cut using an ultra-microtome (Leica Ultracut UCT, Leica Microsystems, Vienna, Austria) and placed on 2 mm × 1 mm slot grids (Plano, Marburg, Germany). Observations were carried out using a JEM-1400 TEM (Jeol, Tokyo, Japan) at 120 kV and a MegaView III camera (Soft Imaging System, Münster, Germany).

### Pit membrane thickness

Pit membrane thickness was measured on TEM images of transverse stem sections, focusing on interconduit pit membranes in the current year, i.e. the latest growth ring. Each central pit membrane thickness value represented the mean of three central measurements, which were distributed around the centre of the pit membrane to cover possible variation. The occurrence of pit channels in bordered pits served as a rough criterion for axial proximity to the pit centre (Kaack *et al*. 2021). For each species and season 10 to 25 pit membranes were measured on one to three sections. Swollen, fresh pit membranes could be identified based on their granular appearance with a transparent to grey colour, which differed from dark, shrunken pit membranes (Fig. 2). When distinctly shrunken and swollen pit membranes were seen within a single sample, we excluded the shrunken pit membranes in our measurements as these could possibly be due to artificial deformation by sample preparation, and may not reflect pit membrane shrinkage in the field (Kotowska *et al*. 2020; Zhang *et al*. 2020). Living conduits could be distinguished from mature, functional conduits based on the presence of cytoplasmic remnants in their pit borders and a homogeneous brightness of their pit membranes. Immature pit membranes of living conduits were not included in our analyses. Images were analysed using ImageJ (National Institutes of Health, Bethesda, Maryland, USA).

### Electron density of pit membranes

To investigate seasonal changes in the brightness or electron density of interconduit pit membranes, TEM images were taken for all eight species and four seasons, focussing again on the latest growth ring only. Grey values of the largest possible area of a pit membrane and of the nearest pit chamber lumen of the same pit were measured, with values between 0 and 255. All samples were treated equally, and only the deviation of grey values was used, not the absolute value. We excluded artefacts such as scratches, folds of the ultra-thin sections, and electron dense components that could be formed by contamination. Although a constant acceleration voltage of 120 kV and electron flux were used, grey values between samples may have been influenced to some extent by minor variation in the thicknesses of the ultra-thin sections and by adjustments to the condenser settings. To limit these effects as much as possible, an automatic linear adjustment of the brightness and contrast was applied during image acquisition. For this reason, absolute grey values were not used for the analyses. However, since the same structures were always measured at a fixed resolution, the grey value difference between the nearest pit chamber lumen (minuend) and the pit membrane (subtrahend) was determined, which was then defined as the relative electron density of the pit membrane.

### Analysis and Statistics

Statistical analyses were performed using R (version 4.44.1, R Core Team 2024) in RStudio (Boston, USA) and the packages data.table, dplyr, lubridate, onewaytests, tidyr, and tidyverse (Grolemund & Wickham 2011; Dag *et al*. 2018; Wickham *et al*. 2019, 2023, 2024; Barrett *et al*. 2024). To assess the overall difference among seasons and species, analyses of variance were conducted. Robust standard errors were estimated using the sandwich package (vcovHC, type = “HC3”; Zeileis et al. 2020) to account for heteroscedasticity, shown by model diagnostics using the DHARMa package (Hartig 2025). Robust Wald tests were used to evaluate the overall significance of species, season, and their interaction (lmtest package; Zeileis & Hothorn 2002). Linear hypothesis tests were applied to assess whether the set of coefficients for species differed jointly from zero (car package; Fox & Weisberg 2019). Estimated marginal means for each season within species were obtained using the emmeans package (Lenth & Piaskowski 2025). Pairwise contrasts were computed with Bonferroni adjustment, incorporating the robust covariance matrix. Data was plotted using ggplot2, ggplotify and multcomp (Hothorn *et al*. 2008; Wickham 2016; Yu 2023). We tested for a correlation between electron density and pit membrane thickness with and without correcting for season and species by subtracting the mean value per species and season form the original value divided by the standard error (Fig S1).

## Results

The robust Wald test indicated that seasons, species, and their interaction had a significant effect on intervessel pit membrane thickness (F(31, 478) = 35, p < 0.001). We observed a decreasing thickness within the current growth ring from summer to winter and the highest thickness in spring for several species (Fig. 1A), although there was considerable interspecific variation. *Betula pendula*, *Prunus avium*, *Fagus sylvatica*, and *Liriodendron tulipifera* had the lowest pit membrane thicknesses during the inactive seasons, meaning in autumn and winter. *Tilia cordata* and *Carpinus betulus* showed similar, but less pronounced patterns, with the thinnest pit membranes occurring in winter. *Acer pseudoplatanus* had a decreasing pit membrane thickness, from summer to spring in the next year. Moreover, *Corylus avellana* had the lowest pit membrane thickness in summer.

**Fig. 1:**
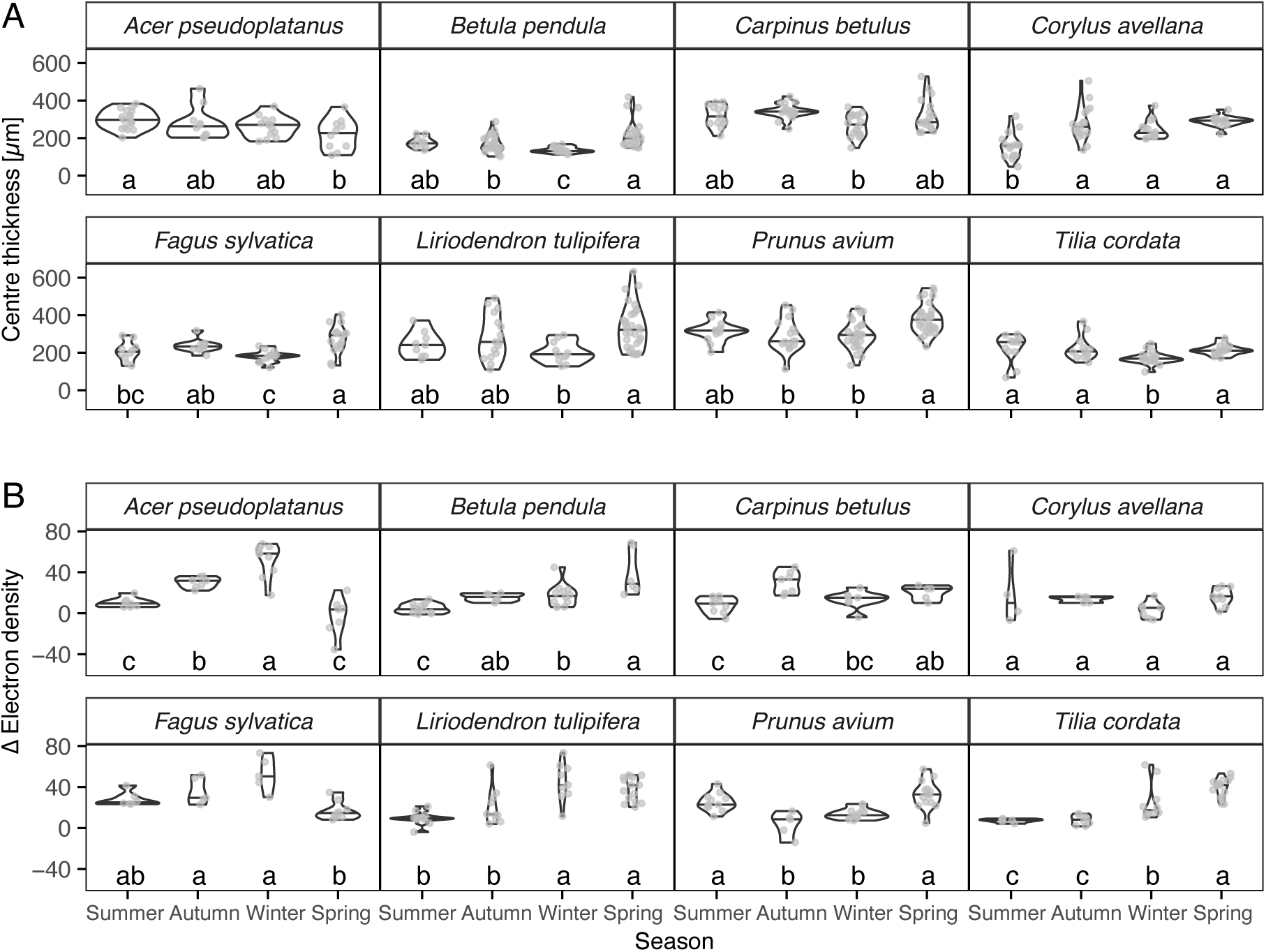
Pit membrane characteristics across species and seasons. A) Pit membrane thicknesses in consecutive seasons. Thickness was measured in the centre of each pit membrane. B) Difference in pit membrane electron density compared to the pit chamber lumen electron density based on the grey values of transmission electron microscopy images. In both plots different letters (a, b, c) indicate statistically significant differences. Violin plots show the distribution of the data and the horizontal line in the plot represents the median.

The seasonal variation in electron density of pit membranes also varied significantly across species (F(31, 240) = 18.23, p < 0.001). At the species level, *Acer pseudoplatanus* and *Fagus sylvatica* had a successively higher electron density from spring to winter (Fig. 1B). *Betula pendula* and *Prunus avium* had an increasing electron density from autumn to spring (Fig. 1B). Here, transparent, pit membranes were limited to vessels that had only recently developed. However, no clear seasonal pattern was observed for *Corylus avellana* and *Carpinus betulus*, while *Liriodendron tulipifera* had a higher electron density in winter and spring than summer and autumn. No correlation was observed between electron density and pit membrane thickness, when data was corrected for species and season (r < 0.001, p = 1.00, 95% CI [–0.03, 0.03])) and only a weak correlation without accounting season and species (r = 0.10, p < 0.001, 95% CI [–0.07, 0.13]); Fig S1).

In general, intervessel pit membranes that become gradually coated, showed a homogeneous increase in electron density, with a granular appearance of the pit membrane. Fairly large particles or deposits, however, were most frequently associated with the outermost layers of intervessel pit membranes, and could not be seen deeper inside pit membranes. When vessel-tracheid pit membranes could be identified under TEM, we observed a pattern of higher electron density on the pit membrane sides that were facing vessel lumina, and a lower electron density on the side of tracheid lumina (Fig. 2). Thus, the diameter of the conduit had to some extent an effect on the seasonal changes in pit membrane coating and electron density.

**Fig. 2:**
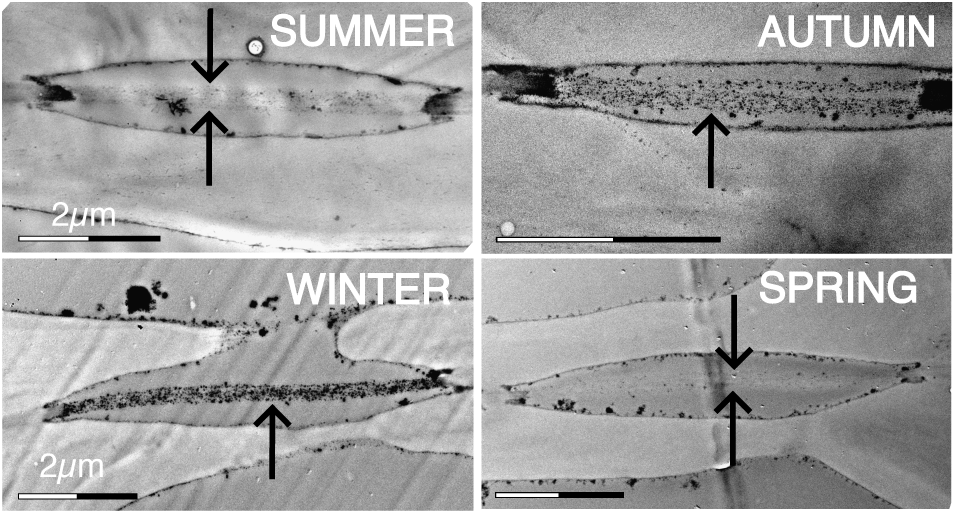
Representative images of pit membranes of *Acer pseudoplatanus* across four consecutive seasons A) summer, B) autumn and C) winter, and D) spring. Pit membranes have a higher electron density by a higher concentration of polar lipids coating the pit membrane.

## Discussion

Despite considerable interspecific variation, our results demonstrate that interconduit pit membranes may undergo changes in pit membrane thickness and electron density across seasons, even within the last growth ring only. For example, seasonality affected both pit membrane thickness and electron density of *Betula pendula* and *Prunus avium*, which is in line with earlier records of shrinkage and coating (Côté & Day 1962; Harada 1962; Wardrop *et al*. 1963; Schmid & Machado 1968). These micromorphological changes within the latest, current year growth ring of stem xylem were remarkable, especially given the challenges to quantify variation in pit membrane thickness and electron density accurately. It is likely that older growth rings show more pronounced deformation of pit membranes compared to those in current year growth rings, as has been reported for instance in sapwood of *Clematis vitalba* (Carmesin *et al*. 2023) and in heartwood of vesselless angiosperms (Zhang *et al*. 2017).

When focusing on the pit membrane thickness, pronounced shrinkage was observed in the species of *Fagus sylvatica*, *Betula pendula*, *Liriodendron tulipifera,* and *Prunus avium*, when the cambium was inactive (Fig. 1A), while pit membrane thickness of *Carpinus betulus* and *Tilia cordata* varied only little among seasons (Seo *et al*. 2007). Based on elasticity measurements, the estimated pit membrane aspiration pressure of *Clematis vitalba* was 2.2 MPa (Carmesin *et al*. 2023), which is a pressure difference highly unlikely to occur between sap-filled vessels in the field. This could also suggest that the shrinkage of pit membranes is unlikely due to pit membrane aspiration caused by the mechanical properties of pit membranes. Pit membrane shrinkage, however, could be caused mechanistically by the hydraulic drag, which is caused by the resistance of the pit membrane to the moving xylem sap, leading also to a gradual decrease in sap velocity over time.

Additionally, it is unclear if shrinkage of pit membranes across bordered pit pairs is symmetric. For instance, shrinkage may happen first on the side of wider conduits than smaller conduits, as a difference in hydraulic drag and also sap flow rate may occur between wide and narrow conduits. In prior studies, interconduit pit membranes that become gradually coated, showed a homogeneous increase in electron density, with a granular appearance of the pit membrane, which we also observed (Figs. 2, S2, S3; Schmid & Machado 1968). In contrast, strong dehydration and shrinkage of pit membranes is known to lead to uniformly dark and thin pit membranes, which also do not have the granular appearance characteristic of fresh pit membranes (Zhang *et al*. 2017; Jansen *et al*. 2018). Fairly large particles or deposits, however, were most frequently associated with the outermost layers of interconduit pit membranes, and could not be seen deeper inside pit membranes. In addition to an increasing electron density from summer to winter and the following spring, we observed a higher electron density on the pit membrane sides that were facing wider conduits, and a lower electron density on the side of smaller conduits (Fig. 3). Thus, conduit diameter explained part of the variation in seasonal changes in pit membrane coating and electron density. The increase in electron density can be caused by polar lipids that coat the inner conduit walls and pit membranes (Liese 1963; Schenk *et al*. 2017, 2018). Based on electron microscopy, these lipids have been shown to be remnants of the cytoplasm of vessel elements (Scott *et al*. 1960; Esau *et al*. 1966). Due to their insoluble nature, these polar lipids either cling onto inner conduit walls, or form micelles, which appear to be pushed with moving xylem sap against intervessel pit membranes, but most likely do not pass them (Guan *et al*. 2022). Since wide vessels should be much more efficient in sap transport than narrow conduits based on the Hagen-Poiseuille law (Sperry *et al*. 2006), the observation that electron density is more pronounced on the larger conduit size is in line with the idea that the lipids move with sap flow. Moreover, the lipid micelles may not easily penetrate a pit membrane, leading to a higher electron density on the wide conduit side than on the narrow conduit size, and a pronounced association of larger lipid droplets with the outermost side of pit membranes (Fig. 3). The increase of lipid deposition on the pit membranes probably reduces the sap flow through the pit membrane, which is also likely to become more compressed by shrinkage. As pit membranes become compressed, their pore volume fraction decreases, with porosity values of pit membranes below 50% in oven-dried samples (Zhang *et al*. 2020). At the same time, this reduced porosity also reduces pore voids, including the smallest pore constriction within a pore pathway across the entire pit membrane thickness. Since the smallest pore constriction determines fluid transport and the dynamics of gas-liquid interfaces (Kaack *et al*. 2021), pit membrane shrinkage is expected to increase embolism resistance. Interestingly, this hypothesis has been confirmed for various species (Sorek *et al*. 2021, 2022; Huang *et al*. 2024; Ma *et al*. 2024). However, the lack of a correlation between electron density and pit membrane thickness (Fig. S1) suggests that an increase in the electron density of pit membranes was probably not directly related to shrinkage and a reduction of the pit membrane porosity.

**Fig. 3:**
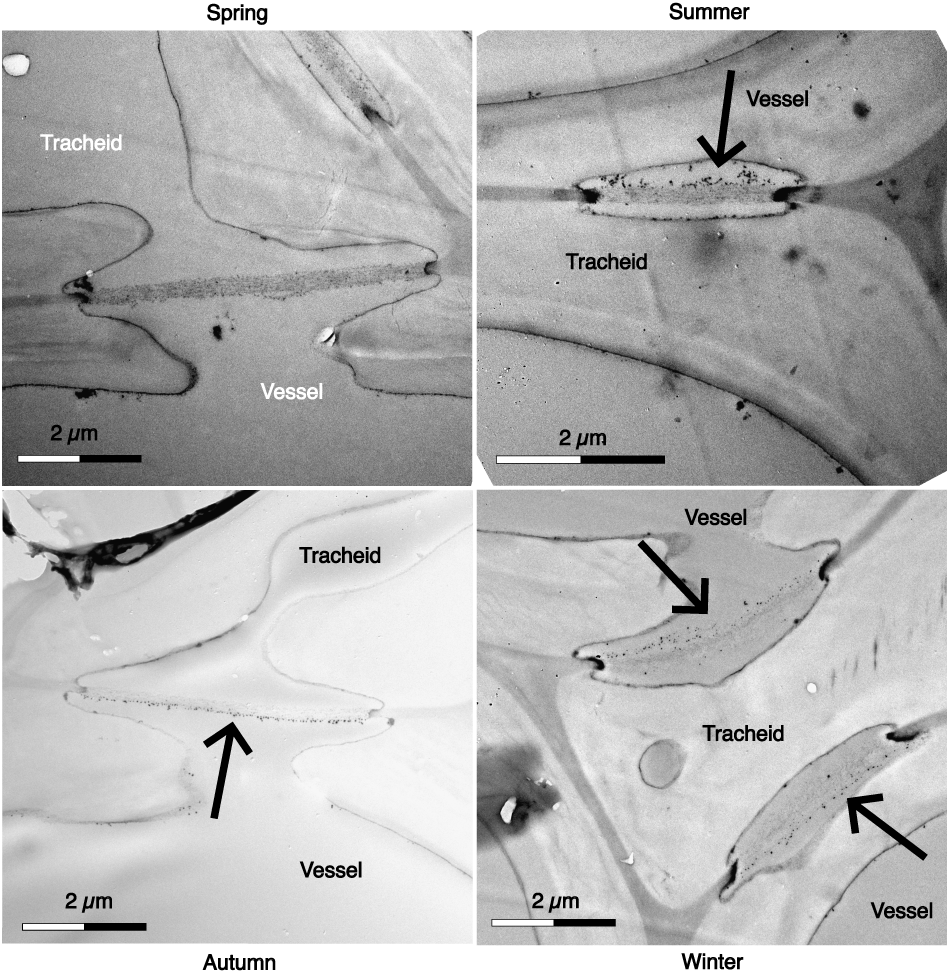
TEM images of pit membranes showing a higher amount of electron dense coating on the side of bigger conduits (probably vessels) than on the side of smaller conduits (probably tracheids). All images are of *Prunus avium*.

Some of the variation in electron density and pit membrane thickness among species might be explained by the different start of cambium activity. Due to differences in angiosperm wood cell kinetics, cambium activity in the species studied starts asynchronously and also depends on temperature and precipitation at the given location, making it difficult to define the exact start of the cambial activity across species (Buttò *et al*. 2019; Noyer *et al*. 2023). To facilitate effective planning and to have the same environmental condition during sampling, seasonal sampling was done around the same time (Table S1). Additional information on summer temperature and precipitation is provided in Fig. S3 and Guan *et al*. (2022). While the cambium of *Acer pseudoplatanus* already becomes active in early spring (Hein *et al*. 2009), *Tilia cordata* and *Liriodendron tulipifera* start growing only when warmer temperatures are reached (Caffarra & Donnelly 2011; LeBlanc *et al*. 2020). This could also explain why *Liriodendron tulipifera* and *Tilia cordata* showed a higher electron density in spring than in summer samples, especially if these species would have started growing later in the year, and the pit membrane measured were not from the same year but from the previous growth season. This could also be the reason why the thinnest pit membranes in *Acer pseudoplatanus* were found in spring (Fig. 1A). The asynchronicity would make it necessary to collect samples of a season not at the same time but to collect samples based on the phenology of the cambium. Although a more frequent sampling strategy would be recommended, we decided in this study to increase the number of species instead of repetitive sampling during the growing season to get a broader species overview.

We observed species dependent patterns of seasonal differences in pit membranes between conduits of several diffuse angiosperms, with pit membrane shrinkage and increased electron density being more pronounced over the course of twelve months in some species than in others. At least part of the intra-tree variation was due to differences between vessels and tracheids, while further research is needed to test how the morphology of pit membranes can be mechanistically linked to the lifespan of conduits and the volume of sapwood in both diffuse porous and ring-porous species.

## Acknowledgements

We thank the Electron Microscopy Section of Ulm University for technical support with preparing TEM-samples and Julia Werner for her support with practical work. SJ acknowledges financial support from the German Research Foundation (DFG, Deutsche Forschungsgemeinschaft; project number 383393940).

## Notes

### Competing Interest Statement

The authors have declared no competing interest.

## References

Barrett T., Dowle M., Srinivasan A., Gorecki J., Chirico M., Hocking T. (2024) data.table: Extension of ‘data.fram’. [online] URL: CRAN.R-project.org/package=data.table (accessed 13 February 2025).

Buttò V., Rossi S., Deslauriers A., Morin H. (2019) Is size an issue of time? Relationship between the duration of xylem development and cell traits. Annals of Botany, 123, 1257–1265.

Caffarra A., Donnelly A. (2011) The ecological significance of phenology in four different tree species: effects of light and temperature on bud burst. International Journal of Biometeorology, 55, 711–721.

Carlquist S. (1975) Ecological strategies of xylem evolution. University of California Press.

Carlquist S. (2001) Comparative wood anatomy: Systematic, ecological, and evolutionary aspects of dicotyledon wood. Springer Berlin Heidelberg.

Carmesin C.F., Port F., Böhringer S., Gottschalk K.-E., Rasche V., Jansen S. (2023) Ageing-induced shrinkage of intervessel pit membranes in xylem of *Clematis vitalba* modifies its mechanical properties as revealed by atomic force microscopy. Frontiers in Plant Science, 14, 1002711.

Catesson A.M. (1983) A cytochemical investigation of the lateral walls of Dianthus vessels. Differentiation and pit-membrane formation. IAWA Journal, 4, 89–101.

Choat B., Cobb A.R., Jansen S. (2008) Structure and function of bordered pits: New discoveries and impacts on whole-plant hydraulic function. New Phytologist, 177, 608– 626.

Christman M.A., Sperry J.S. (2010) Single-vessel flow measurements indicate scalariform perforation plates confer higher flow resistance than previously estimated. Plant, Cell & Environment, 33, 431–443.

Côté W.A.J., Day A.C. (1962) Vestured pits-fine structure and apparent relationship with warts. Technical Association of the Pulp and Paper Industry, 45, 906–910.

Dag O., Dolgun A., Konar N.M. (2018) onewaytests: An R package for one-way tests in independent groups designs. The R Journal, 10, 175–199.

Ellerby D.J., Ennos A.R. (1998) Resistances to fluid flow of model xylem vessels with simple and scalariform perforation plates. Journal of Experimental Botany, 49, 979–985.

Esau K., Cheadle V.I., Gill R.H. (1966) Cytology of differentiating tracheary elements ii. Structures associated with cell surfaces. American Journal of Botany, 53, 765–771.

Fox J., Weisberg S. (2019) An R companion to applied regression, Third. Sage, Thousand Oaks CA. [online] URL: www.john-fox.ca/Companion/ (accessed 13 February 2025).

Fritts H. (1976) Tree rings and climate. Academic Press, London.

Fritts H.C., Vaganov E.A., Sviderskaya I. V, Shashkin A. V (1991) Climatic variation and tree-ring structure in conifers: Empirical and mechanistic models of tree-ring width, number of cells, cell size, cell-wall thickness and wood density. Climate Research, 97–116.

Grolemund G., Wickham H. (2011) Dates and times made easy with lubridate. Journal of Statistical Software 40:1–25. [online] URL: www.jstatsoft.org/v40/i03/ (accessed 13 February 2025).

Guan X., Schenk H.J., Roth M.R., Welti R., Werner J., Kaack L., Trabi C.L., Jansen S. (2022) Nanoparticles are linked to polar lipids in xylem sap of temperate angiosperm species. Tree Physiology, 42, 2003–2019.

Hacke U.G., Sperry J.S., Wheeler J.K., Castro L. (2006) Scaling of angiosperm xylem structure with safety and efficiency. Tree Physiology, 26, 689–701.

Harada H. (1962) Electron microscopy of ultra-thin sections of Beech wood (*Fagus crenata* Blume). Journal of the Japan Wood Research Society, 8, 252–258.

Hartig F. (2025) DHARMa: Residual diagnostics for hierarchical (multi-level / mixed) regression models. [online] URL: github.com/florianhartig/dharma. (accessed 16 December 2025).

Hein S., Collet C., Ammer C., Goff N. Le, Skovsgaard J.P., Savill P. (2009) A review of growth and stand dynamics of *Acer pseudoplatanus* L. in Europe: Implications for silviculture. Forestry: An International Journal of Forest Research, 82, 361–385.

Hothorn T., Bretz F., Westfall P. (2008) Simultaneous inference in general parametric models. Biometrical Journal, 50, 346–363.

Huang R., Di N., Xi B., Yang J., Duan J., Li X., Feng J., Choat B., Tissue D. (2024) Herb hydraulics: Variation and correlation for traits governing drought tolerance and efficiency of water transport. Science of The Total Environment, 907, 168095.

Ingram S., Jansen S., Schenk H.J. (2023) Lipid-coated nanobubbles in plants. Nanomaterials, 13, 1776.

Jansen S., Choat B., Pletsers A. (2009) Morphological variation of intervessel pit membranes and implications to xylem function in angiosperms. American Journal of Botany, 96, 409–419.

Jansen S., Klepsch M., Li S., Kotowska M., Schiele S., Zhang Y., Schenk H. (2018) Challenges in understanding air-seeding in angiosperm xylem. Acta Horticulturae

Kaack L., Altaner C.M., Carmesin C., Diaz A., Holler M., Kranz C., Neusser G., Odstrcil M., Schenk H.J., Schmidt V. (2019) Function and three-dimensional structure of intervessel pit membranes in angiosperms: A review. IAWA Journal, 40, 673–702.

Kaack L., Weber M., Isasa E., Karimi Z., Li S., Pereira L., Trabi C.L., Zhang Y.A., Schenk H.J., Schuldt B. (2021) Pore constrictions in intervessel pit membranes provide a mechanistic explanation for xylem embolism resistance in angiosperms. New Phytologist, 230, 1829–1843.

Kininmonth J.A. (1972) Permeability and Fine Structure of Certain Hardwoods and Effects on Drying. II. Differences in Fine Structure of *Nothofagus fusca* Sapwood and Heartwood. Holzforschung, 26, 32–38.

Kotowska M.M., Thom R., Zhang Y., Schenk H.J., Jansen S. (2020) Within-tree variability and sample storage effects of bordered pit membranes in xylem of *Acer pseudoplatanus*. Trees, 34, 61–71.

LeBlanc D., Maxwell J., Pederson N., Berland A., Mandra T. (2020) Radial growth responses of tulip poplar (Liriodendron tulipifera) to climate in the eastern United States. Ecosphere, 11, e03203.

Lens F., Sperry J.S., Christman M.A., Choat B., Rabaey D., Jansen S. (2011) Testing hypotheses that link wood anatomy to cavitation resistance and hydraulic conductivity in the genus Acer. New Phytologist, 190, 709–723.

Lenth R. V., Piaskowski J. (2025) emmeans: Estimated Marginal Means, aka Least-Squares Means. [online] URL: rvlenth.github.io/emmeans/ (accessed 16 December 2025).

Li S., Lens F., Espino S., Karimi Z., Klepsch M., Schenk H.J., Schmitt M., Schuldt B., Jansen S. (2016) Intervessel pit membrane thickness as a key determinant of embolism resistance in angiosperm xylem. IAWA Journal, 37, 152–171.

Liese W. (1963) Tertiary wall and warty layer in wood cells. Journal of Polymer Science Part C: Polymer Symposia, 2, 213–229.

Ma B.-L., Liao S.-H., Lv Q.-Z., Huang X., Jiang Z.-M., Cai J. (2024) Seasonal plasticity of stem embolism resistance and its potential driving factors in six temperate woody species. Physiologia Plantarum, 176, e14421.

De Micco V., Campelo F., De Luis M., Bräuning A., Grabner M., Battipaglia G., Cherubini P. (2016) Intra-annual density fluctuations in tree rings: How, when, where, and why? IAWA Journal, 37, 232–259.

Noyer E., Stojanović M., Horáček P., Pérez-de-Lis G. (2023) Toward a better understanding of angiosperm xylogenesis: A new method for a cellular approach. New Phytologist, 239, 792–805.

Olson M., Rosell J.A., Martínez-Pérez C., León-Gómez C., Fajardo A., Isnard S., Cervantes-Alcayde M.A., Echeverría A., Figueroa-Abundiz V.A., Segovia-Rivas A. (2020) Xylem vessel-diameter–shoot-length scaling: Ecological significance of porosity types and other traits. Ecological Monographs, 90, e01410.

Pereira L., Flores-Borges D.N.A., Bittencourt P.R.L., Mayer J.L.S., Kiyota E., Araújo P., Jansen S., Freitas R.O., Oliveira R.S., Mazzafera P. (2018) Infrared nanospectroscopy reveals the chemical nature of pit membranes in water-conducting cells of the plant xylem. Plant Physiology, 177, 1629–1638.

Pesacreta T.C., Groom L.H., Rials T.G. (2005) Atomic force microscopy of the intervessel pit membrane in the stem of *Sapium Sebiferum* (Euphorbiaceae). IAWA Journal, 26, 397– 426.

Plavcová L., Hacke U.G., Sperry J.S. (2011) Linking irradiance-induced changes in pit membrane ultrastructure with xylem vulnerability to cavitation. *Plant*, Cell & Environment, 34, 501–513.

Rathgeber C.B.K., Cuny H.E., Fonti P. (2016) Biological basis of tree-ring formation: A crash course. Frontiers in Plant Science, 7

Šako M., Jansen S., Schenk H.J., Netz R.R., Schneck E., Kanduč M. (2026) How lipids suppress cavitation in biological fluids. Journal of Colloid and Interface Science, 703, 139286.

Schenk H.J., Espino S., Rich-Cavazos S.M., Jansen S. (2018) From the sap’s perspective: The nature of vessel surfaces in angiosperm xylem. American Journal of Botany, 105, 172– 185.

Schenk H.J., Espino S., Romo D.M., Nima N., Do A.Y.T., Michaud J.M., Papahadjopoulos-Sternberg B., Yang J., Zuo Y.Y., Steppe K., Jansen S. (2017) Xylem Surfactants Introduce a New Element to the Cohesion-Tension Theory. Plant Physiology, 173, 1177–1196.

Schenk H.J., Michaud J.M., Mocko K., Espino S., Melendres T., Roth M.R., Welti R., Kaack L., Jansen S. (2021) Lipids in xylem sap of woody plants across the angiosperm phylogeny. The Plant Journal, 105, 1477–1494.

Schmid R., Machado R.D. (1968) Pit membranes in hardwoods—Fine structure and development. Protoplasma, 66, 185–204.

Schweingruber F.H. (1996) *Tree rings and environment: Dendroecology.* Paul Haupt AG Bern, Berne.

Scott F.M., Sjaholm V., Bowler E. (1960) Light and electron microscope studies of the primary xylem of *Ricinus communis*. American Journal of Botany, 47, 162.

Seo J.-W., Eckstein D., Schmitt U. (2007) The pinning method: From pinning to data preparation. Dendrochronologia, 25, 79–86.

Silva L.M., Bujnowski B., Pereira L., Miranda M.T., Schenk H.J., Jansen S. (2025) Gas diffusion kinetics in relation to embolism formation and propagation in angiosperm xylem: a mini-review. Acta Horticulturae, 123–134.

Silva L.M., Pereira L., Kaack L., Guan X., Pfaff J., Trabi C.L., Jansen S. (2024) The potential link between gas diffusion and embolism spread in angiosperm xylem: Evidence from flow-centrifuge experiments and modelling. Plant, Cell & Environment, 47, 4977–4991.

Silva L.M., Pfaff J., Pereira L., Miranda M.T., Jansen S. (2024) Embolism propagation does not rely on pressure only: Time-based shifts in xylem vulnerability curves of angiosperms determine the accuracy of the flow-centrifuge method. Tree Physiology, tpae131.

Sorek Y., Greenstein S., Netzer Y., Shtein I., Jansen S., Hochberg U. (2021) An increase in xylem embolism resistance of grapevine leaves during the growing season is coordinated with stomatal regulation, turgor loss point and intervessel pit membranes. New Phytologist, 229, 1955–1969.

Sorek Y., Greenstein S., Hochberg U. (2022) Seasonal adjustment of leaf embolism resistance and its importance for hydraulic safety in deciduous trees. Physiologia Plantarum, 174.

Sperry J.S., Donnelly J.R., Tyree M.T. (1988) A method for measuring hydraulic conductivity and embolism in xylem. Plant, Cell & Environment, 11, 35–40.

Sperry J.S., Hacke U.G., Pittermann J. (2006) Size and function in conifer tracheids and angiosperm vessels. American Journal of Botany, 93, 1490–1500.

Tyree M.T., Zimmermann M.H. (2002) *Xylem structure and the ascent of sap*. Springer Berlin Heidelberg, Berlin, Heidelberg.

Wardrop A.B., Ingle H.D., Davies G.W. (1963) Nature of vestured pits in angiosperms. Nature, 197, 202–203.

Wheeler E. (1981) Intervascular pitting in *Fraxinus americana* L. International Association of Wood Anatomists Bulletin, 2

Wheeler E., Thomas R.J. (1981) Ultrastructural characteristics of mature wood of southern red oak (*Quercus Falcata* michx.) and white oak (*Quercus alba* L.). Wood and Fiber, 13, 169–181.

Wickham H. (2016) ggplot2: Elegant Graphics for Data Analysis. [online] URL: https://ggplot2.tidyverse.org

Wickham H., Averick M., Bryan J., Chang W., McGowan L.D., François R., Grolemund G., Hayes A., Henry L., Hester J., Kuhn M., Pedersen T.L., Miller E., Bache S.M., Müller K., Ooms J., Robinson D., Seidel D.P., Spinu V., Takahashi K., Vaughan D., Wilke C., Woo K., Yutani H. (2019) Welcome to the tidyverse. Journal of Open Source Software, 4, 1686.

Wickham H., François R., Henry L., Müller K., Vaughan D. (2023) dplyr: A grammar of data manipulation. [online] URL: CRAN.R-project.org/package=dplyr (accessed 13 February 2025).

Wickham H., Vaughan D., Girlich M. (2024) tidyr: Tidy messy data. [online] URL: CRAN.R-project.org/package=tidyr (accessed 13 February 2025).

Yamagishi S., Kojima M., Kuroda K., Abe H., Sano Y. (2024b) Seasonal variation of vessel pits in sapwood: Microscopical analyses of the morphology and chemical components of pit membrane encrustations in *Fraxinus mandschurica*. Annals of Botany, 134, 561– 576.

Yu G. (2023) ggplotify: Convert plot to “grob” or “ggplot” object. [online] URL: CRAN.R-project.org/package=ggplotify (accessed 13 February 2025).

Zeileis A., Horthon T. (2002) Diagnostic checking in regression relationships. R News, 3, 7–10.

Zeileis A., Köll S., Graham N. (2020) Various versatile variances: An object-oriented implementation of clustered covariances in *R*. Journal of Statistical Software, 95

Zhang Y., Carmesin C., Kaack L., Klepsch M.M., Kotowska M., Matei T., Schenk H.J., Weber M., Walther P., Schmidt V. (2020) High porosity with tiny pore constrictions and unbending pathways characterize the 3D structure of intervessel pit membranes in angiosperm xylem. *Plant*, Cell & Environment, 43, 116–130.

Zhang Y., Klepsch M., Jansen S. (2017) Bordered pits in xylem of vesselless angiosperms and their possible misinterpretation as perforation plates. Plant, Cell & Environment, 40, 2133–2146.

Zhang Y., Pereira L., Kaack L., Liu J., Jansen S. (2024) Gold perfusion experiments support the multi-layered, mesoporous nature of intervessel pit membranes in angiosperm xylem. New Phytologist, 242, 493–506.

Ziemińska K. (2023) The role of imperforate tracheary elements and narrow vessels in wood capacitance of angiosperm trees. IAWA Journal, 44, 495–508.

Zimmermann M.H. (1983) Xylem structure and the ascent of sap. Springer-Verlag, Berlin.

Zimmermann M.H., Brown C.L. (1971) Trees: Structure and function. Springer-Verlag, New York, Berlin, Heidelberg.

