## Supplementary Information for "Interconduit pit membranes of temperate angiosperms undergo changes in pit membrane thickness and electron density within the final growth ring"

Schepers, J.R., Silva L.M., Carmesin, C., Huppenberger A., Kaack L., Koraim N., Trabi C.L., Jansen S.

### Supplementary Tables

Table S1: Sampling dates of branches for pit membrane thickness and electron density measurements

| Species | Summer | Autumn | Winter | Spring |
|---|---|---|---|---|
| *Acer pseudoplatanus* | 12.08.2019 | 19.11.2019 | 03.02.20 | 23.6.2020 |
| *Betula pendula* | 06.08.2019 | 19.11.2019 | 03.02.20 | 23.6.2020 |
| *Carpinus betulus* | 06.08.2019 | 19.11.2019 | 03.02.20 | 23.6.2020 |
| *Corylus avellana* | 06.08.2019 | 19.11.2019 | 03.02.20 | 23.6.2020 |
| *Fagus sylvatica* | 12.08.2019 | 19.11.2019 | 03.02.20 | 2.6.2020, 23.6.2020 |
| *Liriodendron tulipifera* | 06.08.2019 | 19.11.2019 | 03.02.20 | 23.6.2020 |
| *Prunus avium* | 06.08.2019 | 19.11.2019 | 03.02.20 | 23.6.2020 |
| *Tilia cordata* | 06.08.2019 | 19.11.2019 | 03.02.20 | 23.6.2020 |

Table S2: Range of pit membrane thickness measurements. Mean and standard deviation of measured pit membrane thickness per species in µm

| Species | Range |
|---|---|
| *Acer pseudoplatanus* | 0.27 ± 0.07 |
| *Betula pendula* | 0.18 ± 0.06 |
| *Carpinus betulus* | 0.31 ± 0.07 |
| *Corylus avellana* | 0.24 ± 0.08 |
| *Fagus sylvatica* | 0.23 ± 0.06 |
| *Liriodendron tulipifera* | 0.29 ± 0.11 |
| *Prunus avium* | 0.33 ± 0.09 |
| *Tilia cordata* | 0.21 ± 0.05 |

**Supplementary Figures**

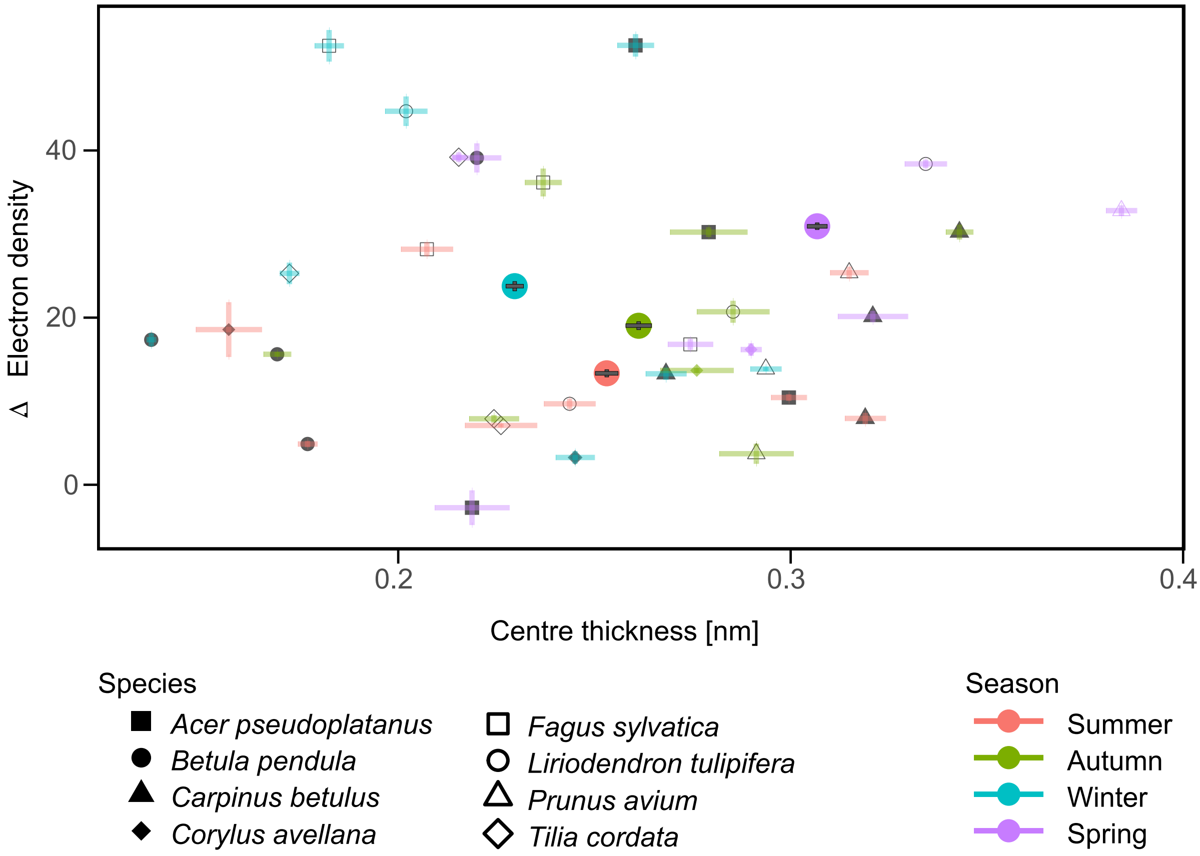


Fig. S1: The pit membrane thickness is plotted on the x-axis versus the relative electron density of pit membranes on the y-axis. The grey symbols show the mean value per species and season with coloured whiskers represent the standard error. Big coloured circles indicate the mean per season with grey wishers showing the standard error per season across all species.

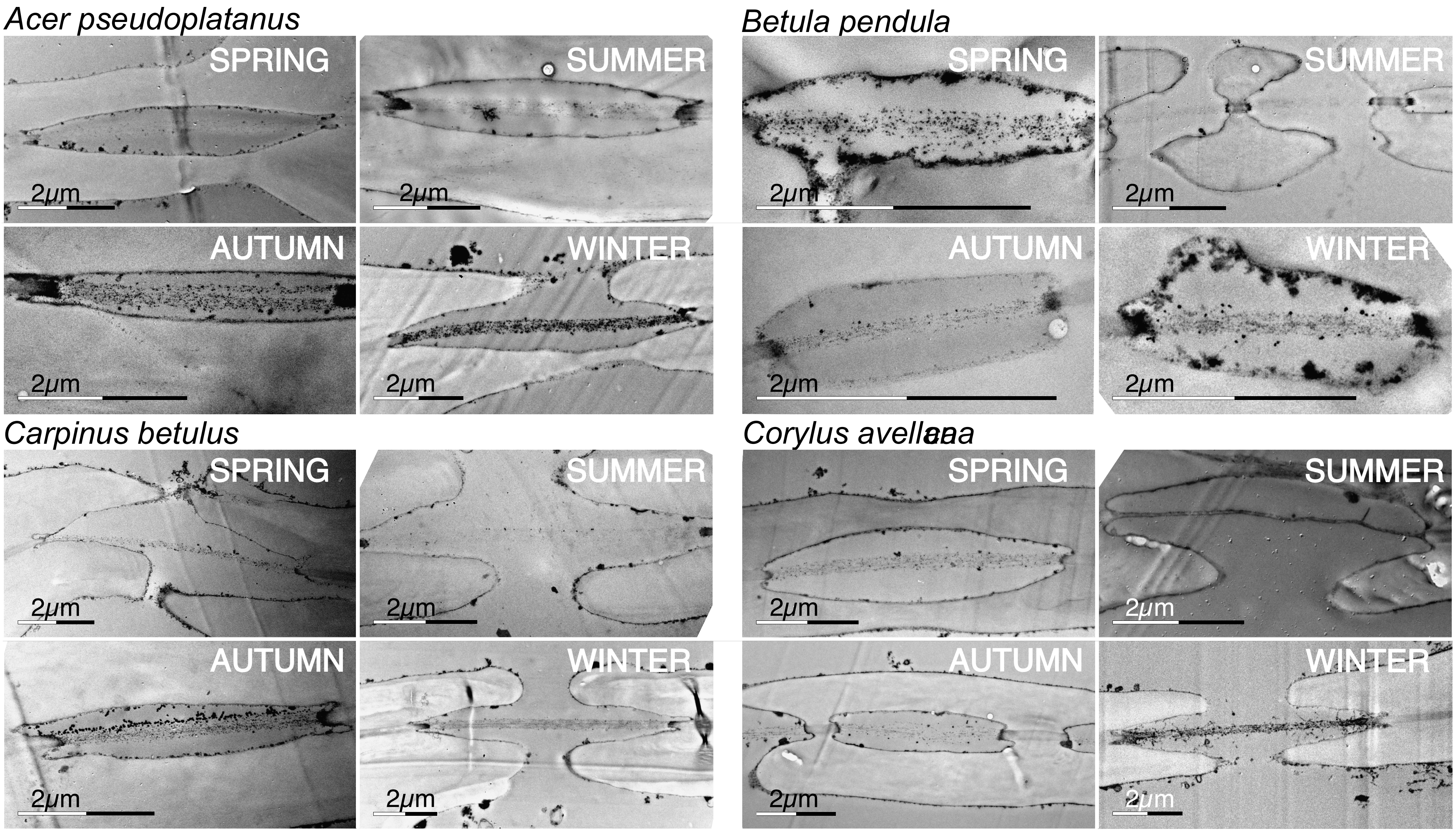


Fig. S2: Representative images of pit membrane of all studied species across the four seasons. Pit membranes appear darker, if there is a higher electron density caused by a higher concentration of polar lipids coating the membrane.

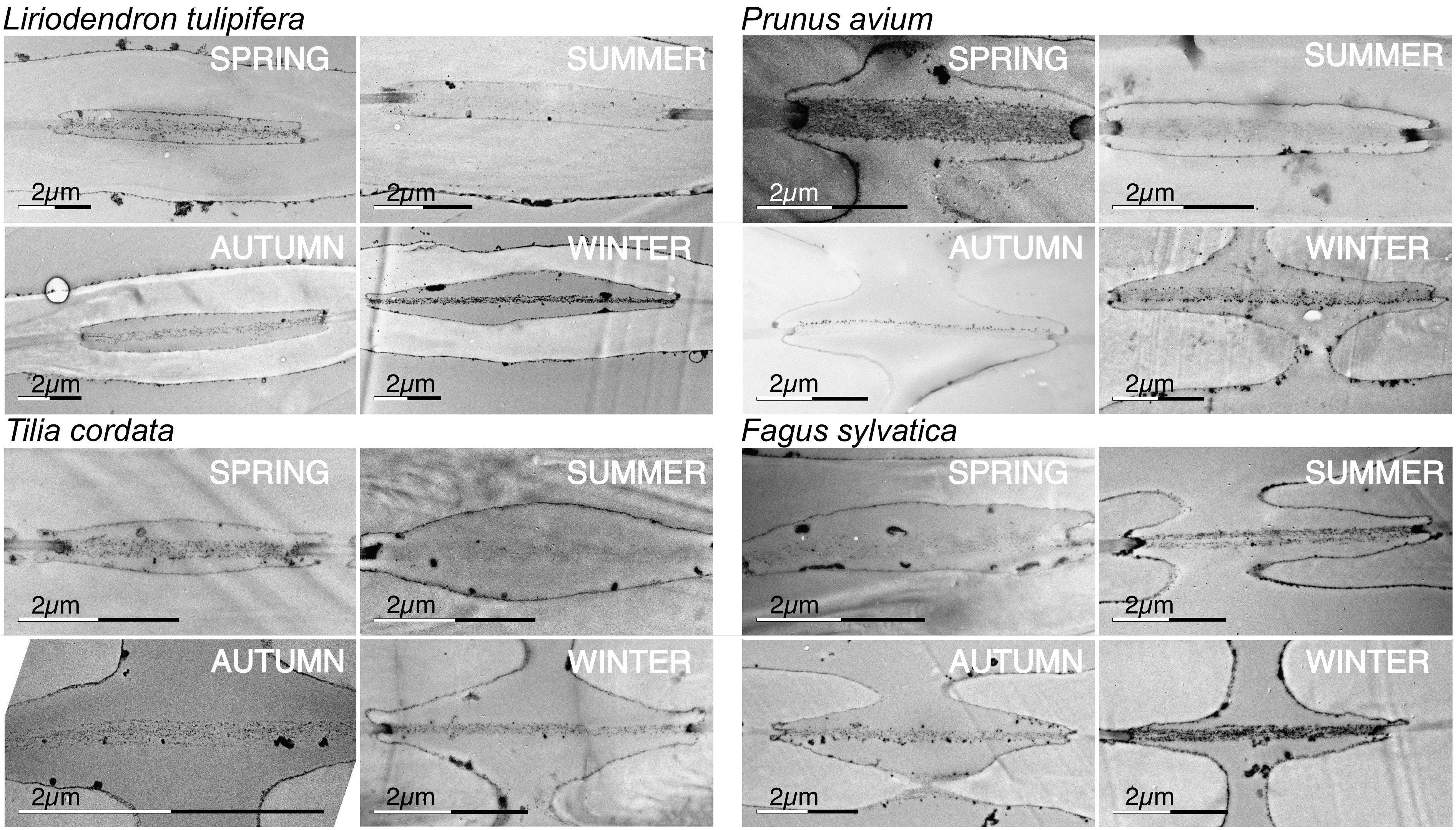


Fig. S2: Representative images of pit membrane of all studied species across the four seasons. Pit membranes appear darker, if there is a higher electron density caused by a higher concentration of polar lipids coating the membrane.

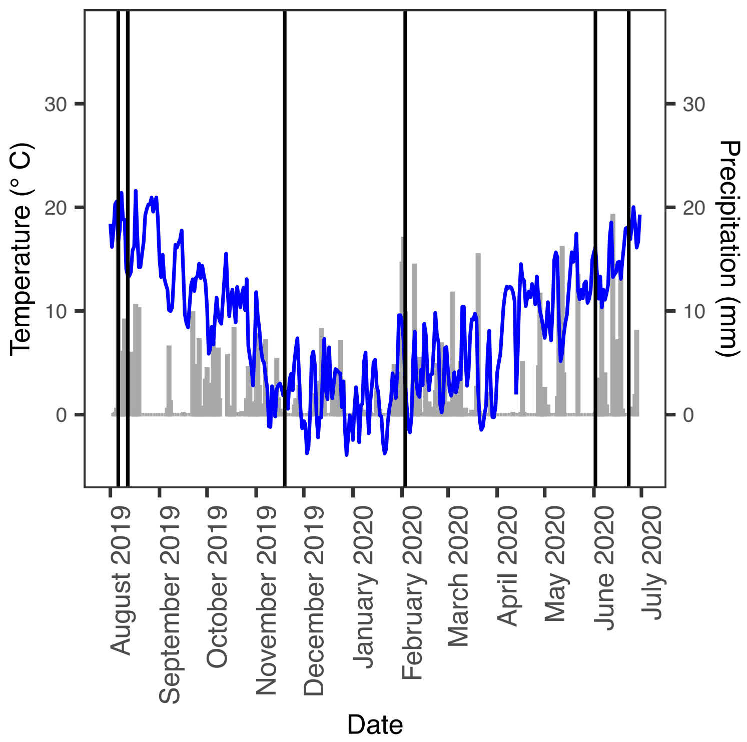


Fig. S3: Daily variations in mean temperature (°C, blue line) and precipitation (mm, grey bars) in summer 2019 to summer 2020. Black vertical dashes indicated dates when samples for pit membrane thickness and electron density measurements were collected. Climate data from Mähringen, Ulm, ca. 2 km from the field site, were obtained from the German weather Service (cdc.dwd.de/portal/)
